# Plasmids, Prophages and Defense Systems are Depleted from Plant Microbiota Genomes

**DOI:** 10.1101/2024.11.17.623994

**Authors:** Avi Bograd, Yaara Oppenheimer-Shaanan, Asaf Levy

**Affiliations:** The Department of Microbiology and Plant Pathology, The Institute of Environmental Science, The Robert H. Smith Faculty of Agriculture, Food & Environment, The Hebrew University of Jerusalem, Rehovot, 76100001, Israel

**Keywords:** Plant-associated bacteria, mobile genetic elements, mobilome, prophages, plasmids, transposons, defense systems, phyllosphere, comparative genomics, metagenomics, microbial ecology

## Abstract

Plant-associated bacteria significantly impact plant growth and health. Understanding how bacterial genomes adapt to plants can provide insights into their growth promotion and virulence functions. Here, we compared 38,912 bacterial genomes and 6,073 metagenomes to explore the distribution of mobile genetic elements and defense systems in plant-associated bacteria. We reveal a consistent taxon-independent depletion of prophages, plasmids, and defense systems in plant-associated bacteria, particularly in the phyllosphere, compared to other ecosystems. The mobilome depletion suggests the presence of unique ecological constraints or molecular mechanisms exerted by plants to control the bacterial mobilomes independently of the bacterial defense.

## Background

The microbial world is a driving force behind the evolution of life on Earth, with bacteria playing a pivotal role in shaping ecosystems through their interactions with other organisms and the environment [1]. Mobile genetic elements (MGE) such as plasmids, prophages, and transposons are key agents of genetic and evolutionary change, enabling horizontal gene transfer (HGT) and adaptation to environmental pressures [2]. However, these elements often impose fitness costs on their hosts, like resource consumption and potential disruption of vital genes. In order to combat MGE, bacteria have evolved defense systems such as restriction enzymes, CRISPR-Cas, abortive infection, and many new systems [3–7].

Several studies revealed habitat-specific differences in MGE distribution. For example, soil encodes much more plasmid taxonomic units than humans [8]. Similarly, phages show habitat specificity, with some restricted to particular environments [9]. However, comprehensive studies on the abundance of MGE and defense systems in plant-associated (PA) environments are scarce. PA environments present unique challenges, including fluctuating nutrients, abiotic stresses, microbial competition, and bacteria-plant immunity interactions. While some plasmids in PA bacteria, like rhizobial nodulation genes and Agrobacterium Ti plasmids, are thoroughly studied, the overall distribution of MGE and their associated defense systems in PA bacteria remains poorly understood. Only a few works study phages in the plant environment beyond a biocontrol against plant pathogens [10–14]. Our previous work identified MGE depletion in PA bacteria, but it was limited in scope and did not include defense systems [15]. Here, we perform a large-scale comparative genomics and metagenomics analysis to explore the distribution of MGEs and defense systems in PA bacteria, addressing this knowledge gap.

## Results and Discussion

We conducted a comprehensive analysis of 38,912 bacterial isolate genomes (Supplementary Table 1) and 6,073 shotgun metagenomes (Supplementary Table 2) to explore the distribution of MGE and defense systems in PA bacteria. The isolates were derived from 19 bacterial families across four major phyla: Proteobacteria, Firmicutes, Actinobacteria, and Bacteroidetes. These genomes were categorized into PA, non-plant associated (NPA), or soil bacteria based on their isolation sites. Each genome was annotated for protein domains linked to prophages, plasmids, transposons, and defense systems using a carefully curated list of ∼800 Pfam domains [16] (Supplementary Table 3). The abundance of each domain category was normalized by calculating the relative fraction of each domain category by the total number of domains identified in each genome. This approach enabled a functional comparison of domain distributions across different bacterial families and environments.

Our analysis revealed a consistent and significant depletion of all three MGE classes, plasmids, prophages, and transposons, in PA bacteria compared to NPA bacteria (Figure 1a). Specifically, in 30 out of 57 possible family-MGE class comparisons, NPA bacteria exhibited a higher abundance of MGE than PA bacteria, while the remaining 27 comparisons showed no significant difference between the groups. Remarkably, there were no instances where PA bacteria had a higher abundance of MGEs than their NPA counterparts, demonstrating a clear trend that spans across taxa. This effect was observed in all four major bacterial phyla, suggesting that the MGE depletion is a broad, taxonomy-independent phenomenon. Notably, six families (Microbacteriaceae, Micrococcaceae, Micromonosporaceae, Weeksellaceae, Enterobacteriaceae, and Pseudomonadaceae) showed a depletion of all three MGE classes in PA bacteria, emphasizing the robustness and consistency of this finding.

**Figure 1:**
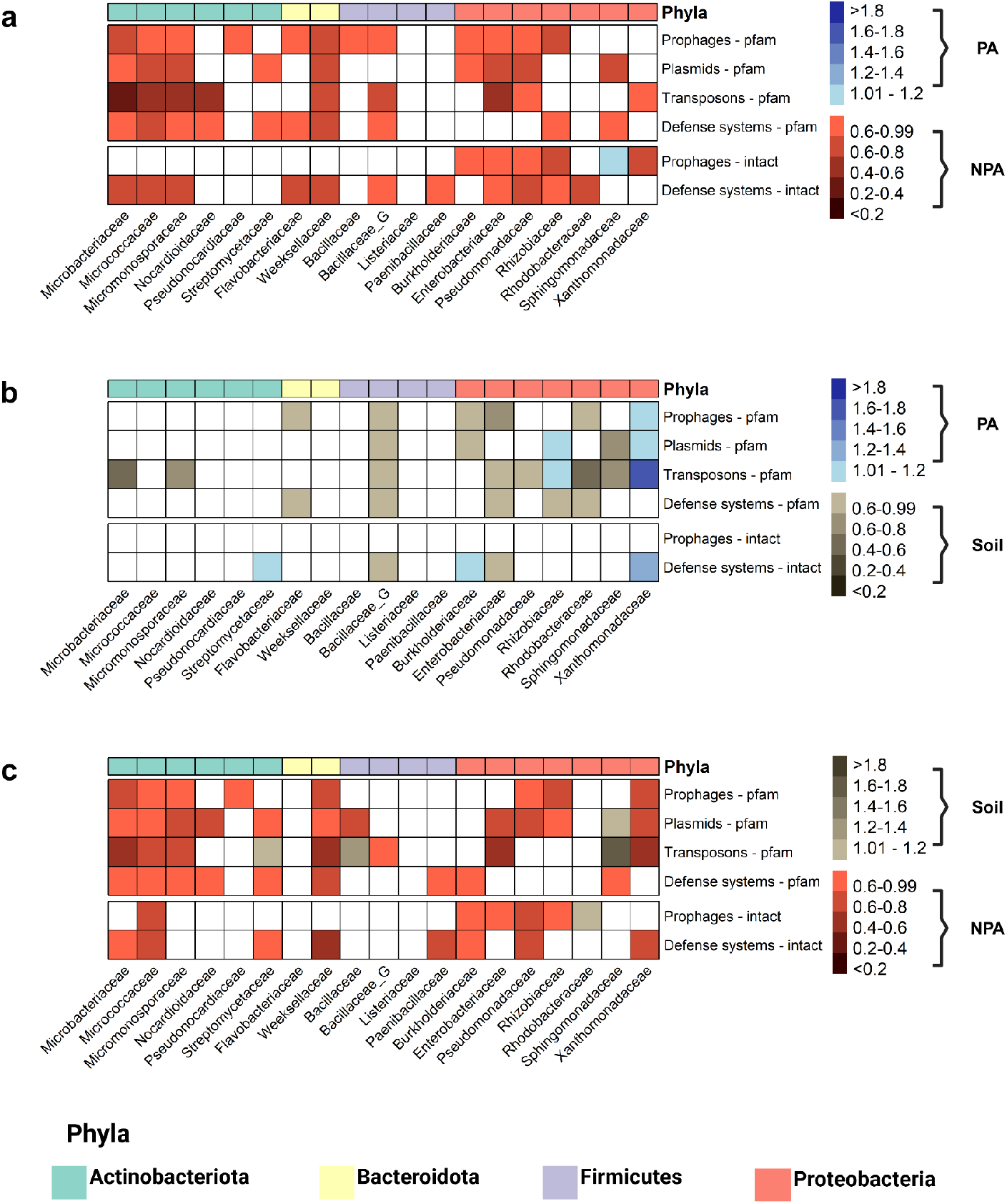
Mobile genetic elements and defense systems are depleted from plant-associated bacteria independently of taxa and MGE class. The abundance of protein domains that belong to MGEs and defense systems as well as the distribution of intact prophages and defense systems were compared across different bacterial families and three types of bacteria: PA bacteria, soil derived bacteria (soil), and NPA bacteria. Each cell represents the fold change of a specific functional group (category) within a bacterial family between two different habitats. The fold change is calculated by dividing the median normalized abundance in one habitat (e.g., PA) by the median normalized abundance in the other habitat (e.g., NPA). **a. Comparison between PA and NPA bacteria, b. Comparison between PA and soil bacteria, c. Comparison between NPA and soil bacteria**. Color coding for abundance comparisons: Blue shades: Higher abundance in PA bacteria. Green shades: Higher abundance in Soil bacteria. Red shades: Higher abundance in NPA bacteria. White: No significant difference between the groups (Wilcoxon–Mann–Whitney test p - value > 0.05).

One possible explanation for the observed depletion of MGEs in PA bacteria is the enrichment of bacterial defense systems in these environments, which might prevent the transfer, integration or genomic persistence of MGE. To investigate this hypothesis, we conducted a similar analysis focusing on the abundance of defense systems in PA and NPA bacteria (Figure 1a). Contrary to the hypothesis, we found that defense systems were also depleted in PA bacteria. Out of the 19 taxonomic families examined, ten displayed significant depletion of defense systems in PA bacteria, while the remaining nine showed no significant difference. This pattern suggests that the observed MGE depletion in PA bacteria cannot be explained by the presence of defense systems. Instead, it points toward a different ecological or molecular mechanism underlying the reduced prevalence of MGEs in PA environments.

To further explore the ecological factors driving this MGE depletion, we compared the relative abundance of MGE and defense systems in PA bacteria with those in soil bacteria (Figure 1b). Soil serves as a reservoir for many PA bacteria, particularly those found in the rhizosphere [17,18]. PA bacteria generally exhibited either a lower or similar abundance of MGE compared to soil bacteria, suggesting that MGE depletion is primarily a PA effect rather than a general soil-related phenomenon. The distinct reduction of MGE in PA bacteria implies that the plant environment imposes unique selective pressures that are not present in soil alone. When comparing soil to NPA bacteria (Figure 1c), we observed a trend where soil bacteria had lower or similar MGE ratios than NPA, suggesting that MGE depletion in PA bacteria is affected by the selection acting against these in both soil and plant environments. This pattern was consistent across different MGE classes and taxa, highlighting the plant-specific effects on the bacterial mobilome.

An interesting exception to this trend was observed in the Xanthomonadaceae family, where soil bacteria showed a lower fraction of MGEs than both PA and NPA bacteria (Figure 1b-c). This unique pattern in Xanthomonadaceae, a family of mostly plant pathogens [19], suggests that the relationship between MGE abundance and plant association may be more complex in certain bacterial lineages, possibly related to pathogenic lifestyles.

We repeated the pfam-based analysis with intact prophages [20,21] and intact defense systems [22]. The analysis largely corroborated the Pfam domain analysis, with some nuances (Figure 1). For intact prophages, the depletion effect was particularly pronounced within the Proteobacteria phylum. In the comparison between PA and NPA bacteria, five out of seven Proteobacteria families exhibited significant depletion in PA bacteria. Interestingly, the comparison between PA and soil bacteria revealed no significant differences across all families, suggesting that the soil environment may play a role in shaping prophage abundance in PA bacteria. Regarding intact defense systems, we observed a widespread depletion among PA bacteria across bacterial families. In the PA versus NPA comparison, 11 out of 19 families exhibited significant depletion in PA bacteria, with the remaining families showing no significant difference. The comparison between PA and soil bacteria revealed a more complex pattern, with three families (Streptomycetaceae, Burkholderiaceae, and Xanthomonadaceae) showing enrichment in PA bacteria, while two families (Bacillaceae_G and Enterobacteriaceae) demonstrated enrichment in soil bacteria. The intact defense systems findings further support the overall trend of MGE and defense system depletion in PA bacteria, while also highlighting some taxon-specific variations in this pattern.

To validate and extend our findings from isolate genomes, we conducted a comparative metagenomics analysis of 6,073 shotgun metagenomes from diverse environmental habitats (Figure 2). This analysis not only confirmed our isolate genome findings but also provided a more detailed view of MGE and defense system distribution across different environments. PA metagenomes, particularly those from the phyllosphere, consistently exhibited the lowest normalized abundance of prophages, plasmids, and defense systems among all habitats examined (Figure 2a,b,d). This trend was especially pronounced when comparing PA metagenomes to those from mammals, insects, fungi, and fish. Intriguingly, even green and red algae showed higher abundance compared to land plants, despite their shared evolutionary ancestry. This suggests that the depletion of MGE and defense systems is a specific adaptation to terrestrial plant environments rather than a general feature of photosynthetic hosts.

**Figure 2:**
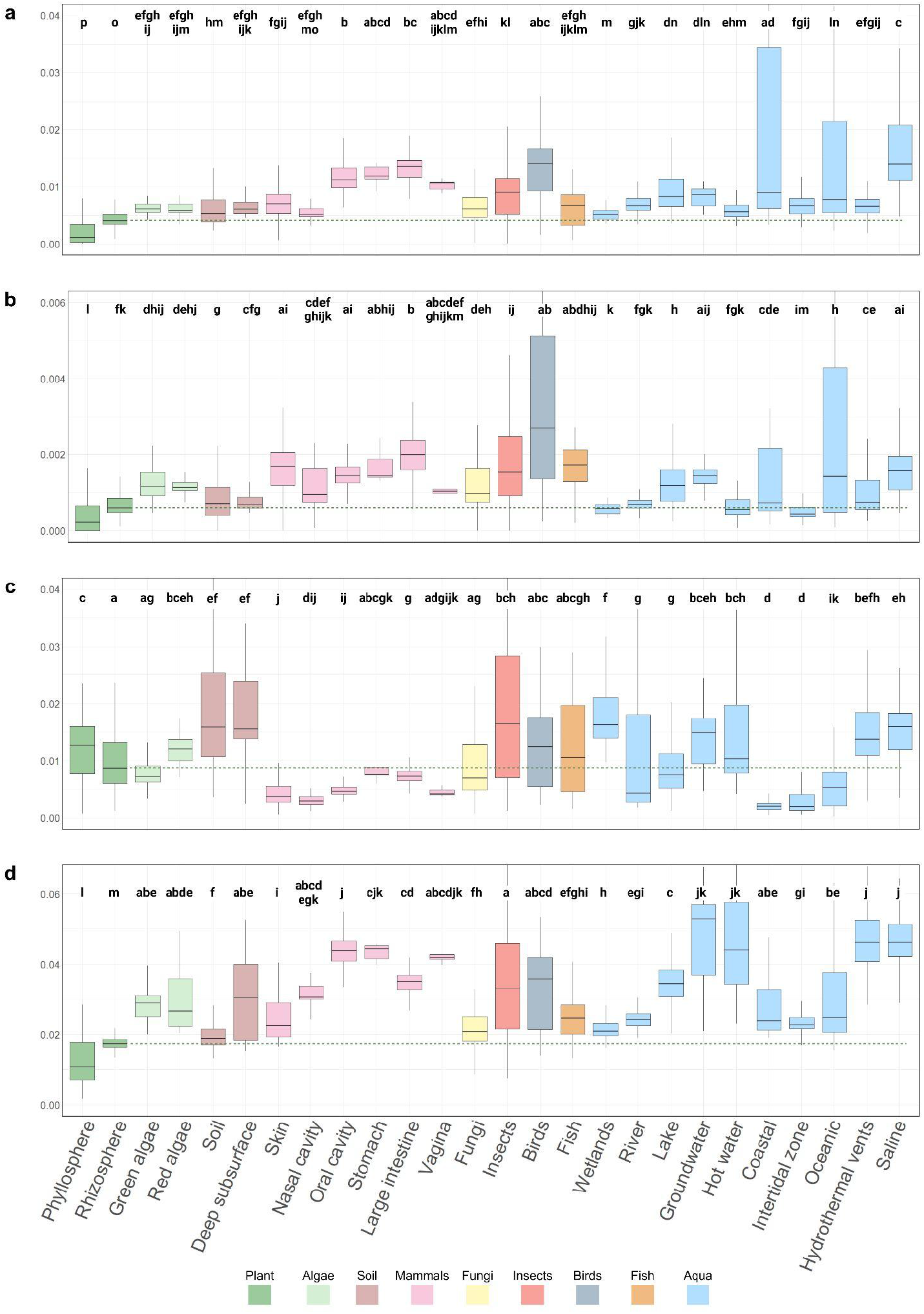
Distribution of Mobile Genetic Elements Across Different Habitats. The figure presents box plots showing the normalized abundance of four types of MGE across various habitats. Each panel (a-d) represents a different MGE category: a) Prophages b) Plasmids c) Transposons d) Defense systems. The x-axis in each panel represents different habitats, including plant-associated environments (e.g., phyllosphere, roots), soil, marine, and various host-associated environments. The y-axis shows the normalized abundance of Pfam domains associated with each MGE type. Letters above the boxes indicate statistically significant differences between habitats (Kruskal-Wallis p-value < 0.05 and dunn test adj p - value < 0.01 (habitats sharing the same letter are not significantly different)).

Within PA metagenomes, the depletion trend was particularly strong in phyllosphere samples, which consistently exhibited the lowest normalized abundance of these elements among all habitats examined. While exposure to fluctuating environmental conditions and low nutrient availability likely play a role, these factors alone cannot fully explain the observed pattern. Other habitats that are subjected to similar ecological pressures, such as mammalian skin or coastal marine environments, do not show the same trend. For instance, our analysis revealed that the microbiome of the mammalian skin and coastal zones, which are also exposed to UV radiation, temperature fluctuations, and variable moisture levels, maintain higher levels of MGE and defense systems compared to the phyllosphere. This suggests that unique aspects of the plant environment, beyond general ecological stressors, contribute significantly to the observed depletion. Possible plant-specific factors could include the chemical composition of plant exudates, specific plant immune responses, or the physical structure of plant surfaces. Another possible factor is wet-dry cycle which is common in the phyllosphere and confers antibiotic resistance [23]. The fact that the MGE depletion is observed not only in the phyllosphere but also, to a lesser extent, in the rhizosphere, which is not directly exposed to many of these external stressors, further supports the hypothesis that plant-specific factors play a crucial role in shaping the mobilome of their associated bacterial communities.

Interestingly, transposons in metagenomes did not follow the same depletion pattern observed for other MGE (Figure 2c). Unlike prophages and plasmids, transposon-related domains did not show a significant reduction in PA metagenomes compared to other ecosystems. Instead, there was a notable decrease in transposons within mammalian-associated habitats, which belong to the NPA category. This divergence suggests that transposons might play a unique role in PA bacterial communities, possibly facilitating rapid adaptation to the dynamic plant environment. The retention of transposons (according to the metagenome but not the isolate genomes analysis), warrants further investigation into their specific functions and selective advantages in plant-microbe interactions. One clear distinction between transposons, plasmids and prophages is the extracellular phase of the two latter MGE groups suggesting an environmental effect leading to the MGE depletion.

Our results indicate that PA environments impose unique ecological pressures on their bacterial inhabitants, thereby reshaping their mobile gene pool. One possible explanation is that specific interactions between PA bacteria and plant cells could create conditions that disfavor the retention of MGE. For example, the low bacterial cell density typically found in the phyllosphere [24] could limit the likelihood of MGE exchange, further relaxing selection pressures for maintaining MGE and defense systems. The fact that prophages are underrepresented in plant microbiota suggests that phages in general are relatively scarce in this environment, likely due to low bacterial density (in the phyllosphere and endosphere) or non-specific antiviral compounds released by plants. Namely, we propose an hypothesis that plant immunity may reduce MGEs in their bacterial communities, reducing the need for bacteria to maintain their genetically encoded defense systems. If indeed bacteria are protected from phages by plant-based mechanisms, then plant hosts may partially serve as “cities of refuge” for bacteria (a biblical term for places where individuals could request for asylum from their attackers), in some sense. Future studies should measure cell-free phage titers in the plant environment compared to other ecosystems and test for antiphage activity by plant compounds.

These findings have broad implications for our understanding of plant-microbe interactions and bacterial genome evolution. The depletion of MGE and defense systems in PA bacteria could impact the evolution of antibiotic and metal resistance, as MGE often carry resistance genes. Furthermore, the reduced genomic plasticity in PA bacterial communities may influence their adaptability to environmental changes, such as shifts in agricultural practices or climate conditions. This depletion could also affect the ability of PA bacteria to acquire beneficial traits through horizontal gene transfer, potentially shaping the co-evolutionary dynamics between plants and their associated microbiota.

Despite these insights, our interpretation requires careful consideration of potential caveats. The plant environment may harbor novel MGE and defense systems not detected by our current methods, which rely on known genetic elements. This limitation could be addressed in future studies employing more sensitive detection techniques, such as functional metagenomics, to capture a broader diversity of MGE. Additionally, variations in MGE and defense system abundance among different plant species need further exploration. Detailed studies focusing on specific plant-microbe systems could reveal species-specific mechanisms driving the observed depletion, with important implications for agriculture and ecosystem management.

## Conclusions

Our study demonstrates a consistent depletion of prophages, plasmids, and defense systems in PA bacteria, particularly in the phyllosphere, across diverse bacterial taxa which is supported by both cultured and non-cultured bacteria. This mobilome depletion, likely driven by both plant-specific and soil-specific factors, represents a fundamental shift in the genomic landscape of PA microbes. These findings reveal a previously unrecognized aspect of plant-microbe interactions, suggesting that plants may exert unique selective pressures on their microbiota, effectively controlling bacterial genome composition by a yet unknown mechanism. Our study opens new avenues for research into the mechanisms shaping evolution of plant-associated bacterial communities and their functional consequences.

## Methods

### Data Compilation and Filtering

A total of 38,912 isolate bacterial genomes were analyzed, sourced from 19 phylogenetic families across four different phyla, and categorized into PA, NPA, and soil habitats based on their isolation sites (Supplementary Table 1). The dataset was derived from public data sources [15,25–32]. Genomes were filtered for completion (>95%), contamination (<5%), and N50 scores (>50,000 bp).

A dataset of 6,073 metagenomes representing diverse environments was obtained from the IMG database [27] (Supplementary Table 2). Metagenomes with fewer than 10,000 Pfam domains were excluded to ensure robust analysis.

### Pfam Domain Classification

Four distinct Pfam [16] lists were compiled, each related to prophages (n = 322), plasmids (n = 81), transposons (n = 102), and bacterial defense systems (n = 285). Transposon and prophage Pfams were identified using keyword searches in the InterPro database [33].

Plasmid-associated Pfams were obtained from Deeplasmid [34] and IMG-PR databases [8]. Defense system Pfams were compiled from published studies [3,35]. The complete list of Pfam domains for each category is provided in (Supplementary Table 3).

### Prophage Identification and Validation

Intact prophages were identified within bacterial genomes using VirSorter2 [20] with parameters set to include dsDNA and ssDNA phages, retaining the original sequence. Only sequences with a minimum length of 5,000 bp and a VirSorter score of at least 0.5 were included in the analysis.

To validate the prophage predictions, CheckV was employed [21]. The validation process involved using the final viral sequences generated by VirSorter2, which were then analyzed by CheckV to assess genome completeness. Only prophage sequences classified by CheckV as “provirus” and with a quality classification of “Medium” or higher (indicating at least 80% completeness) were retained for further analysis.

For each bacterial genome, the total proviral length and gene counts, including both viral and host genes, were summed. The proviral length for each genome was normalized by dividing by the total nucleotide count of the genome (Supplementary Table 4).

### Defense System Identification and Analysis

Defense systems within bacterial genomes were identified using the Defense-finder tool [22]. The tool was run using a custom database to annotate the protein sequences associated with defense systems in each genome. The resulting data were grouped by defense system type for each genome.

For each defense system type, the total protein length (in amino acids) was calculated and converted to base pairs (bp) by multiplying by 3, to account for codon length. The total bp length for each defense system type was normalized by dividing by the total nucleotide count of the corresponding bacterial genome, ensuring consistency across samples (Supplementary Table 5).

### Data Analysis

For each Pfam category, domain occurrences were quantified across all bacterial genomes and metagenomes, with counts normalized by dividing by the total number of Pfam domains identified.

To assess statistical differences among isolate bacteria, intact prophages, and intact defense systems, the Wilcoxon rank-sum test was employed, as it is appropriate for non-normal distributions. Fold changes were calculated for each distinct phylogenetic family by taking the median normalized count in one habitat group and dividing it by the median normalized count in the comparison habitat group.

For metagenomes, the Anderson-Darling test revealed non-normal distribution. The Kruskal-Wallis test was used for group comparisons, followed by Dunn’s test with Benjamini-Hochberg adjustments for post-hoc pairwise comparisons (adjusted p-value < 0.01). The complete results of Dunn’s test, including all pairwise comparisons and adjusted p-values, are presented in Supplementary Table 6.

All statistical analyses and visualizations were performed using R packages: readr (2.1.4) dplyr (1.1.3) tidyr (1.3.0) pheatmap (1.0.12) RColorBrewer (1.1-3) ggplot2 (3.5.0) dunn.test (1.3.5) multcompView (0.1-10) rcompanion (2.4.34). R scripts available at https://github.com/avi-bo/MGEs

## Acknowledgements

AL is generously supported by the Israeli Science Foundation (Grants #1535/20, #3062/20), the Israeli Ministry of Innovation, Science, and Technology (Grant #1001695377), and Israel Innovation Authority (Grant #81259), ICA in Israel, Israel Ministry of Agriculture (Grant 12-12-0008), and the Volkswagen Stiftung (Grant ZN4041).

